# Evaluation of Oxford Nanopore Sequencing for Antimicrobial Resistance Surveillance in *Salmonella*: Comparison with Phenotypic Antimicrobial Susceptibility in a Large-Scale Study

**DOI:** 10.64898/2026.05.19.726213

**Authors:** Yu-Ping Hong, Ying-Shu Liao, You-Wun Wang, Shao-Chun Kuo, Ru-Hsiou Teng, Shiu-Yun Liang, Jui-Hsien Chang, Hsiao Lun Wei, Chien-Shun Chiou

## Abstract

*Salmonella* is a major zoonotic foodborne pathogen, and antimicrobial resistance (AMR) in *Salmonella* presents a significant public health challenge. Whole-genome sequencing (WGS) offers a more rapid and comprehensive method for AMR characterization compared to conventional antimicrobial susceptibility testing (AST), supporting antimicrobial therapy and surveillance efforts. In this study, Oxford Nanopore Technology (ONT)-based WGS was performed on 1,490 *Salmonella* isolates collected through nationwide surveillance in Taiwan in 2025. Genotypic resistance inferred from WGS data was compared with phenotypic AST results to assess the performance of ONT-WGS. Overall, WGS-inferred resistance showed high concordance with phenotypic resistance for most antimicrobials. However, major genotype– phenotype discordance was observed, attributed to four categories: (i) breakpoint-dependent classification, (ii) reduced or absent phenotypic expression of resistance genes, (iii) MIC modulation by *ramAp*, and (iv) absence of known AMR determinants. Notable discrepancies included tigecycline resistance without known genetic determinants, nalidixic acid resistance linked to *ramAp*-mediated MIC elevation, and a high prevalence of colistin resistance (35.4%) in *S*. Enteritidis without identifiable AMR determinants. Additionally, a significant proportion of ESBL- and AmpC-producing isolates were classified as susceptible or intermediate to cefotaxime and ceftazidime under CLSI criteria, highlighting the potential for misclassification and treatment failure. These findings demonstrate that ONT-WGS enables accurate, comprehensive AMR characterization, offering direct identification of AMR determinants and minimizing misclassification due to breakpoint-based AST interpretations. When interpreted appropriately, WGS can support better antimicrobial selection and serve as a valuable alternative to conventional susceptibility testing.

## INTRODUCTION

*Salmonella* is a leading zoonotic foodborne pathogen globally, responsible for millions of cases of gastroenteritis each year. Antimicrobial resistance (AMR) in *Salmonella* has become a critical issue, as it diminishes the effectiveness of treatment options, exacerbating morbidity, mortality, and healthcare costs (1, 2). The World Health Organization has elevated fluoroquinolone-resistant Salmonella to a higher priority, designating it as a pathogen in need of urgent surveillance and control (3). In Taiwan, long-term surveillance has revealed a steady rise in Salmonella resistance to key antimicrobials (4). Over the past decade, high-risk multidrug-resistant (MDR) *Salmonella* clones have repeatedly emerged, causing large-scale outbreaks. Notable examples include MDR *S*. Anatum, azithromycin-resistant extensively drug-resistant (AziR-XDR) *S*. Typhimurium, MDR *S*. Goldcoast, MDR *S*. Agona, MDR *S*. Infantis, and, more recently, AziR-XDR *S*. Kentucky (5-10).

Antimicrobial susceptibility testing (AST) has long been the gold standard for determining phenotypic resistance, but it is constrained by factors such as lengthy turnaround times and a limited spectrum of antimicrobials tested. Although AST can identify resistance even in the absence of known antimicrobial resistance (AMR) determinants, it has limitations when the MIC falls below the breakpoint despite the presence of resistance genes, potentially resulting in treatment failure. In contrast, whole-genome sequencing (WGS) has become a more efficient and comprehensive alternative, as it enables the direct identification of AMR determinants, precise serotype prediction, high-resolution genotyping, and source attribution (11-14). Unlike AST, WGS can detect all known AMR determinants, including those linked to drugs not be included in the AST panels.

Illumina and Oxford Nanopore Technologies (ONT) are the two most widely used platforms for WGS of bacterial isolates. Illumina short-read sequencing is well-established for its high per-base accuracy and robust performance in AMR detection and genomic epidemiology. However, its long turnaround time and lack of flexibility in deployment are limiting factors for real-time AMR detection and disease surveillance. Illumina sequencing is typically high throughput, making it cost-effective only when sequencing large numbers of bacterial isolates simultaneously. In contrast, ONT long-read sequencing offers faster turnaround times, greater flexibility in deployment, and cost-effectiveness for sequencing smaller numbers of isolates, making it more adaptable for bacterial WGS. While earlier ONT platforms faced challenges with basecalling errors (15), recent advances in nanopore chemistry and basecalling algorithms have significantly improved accuracy, with studies showing comparable performance to Illumina for core-genome multilocus sequence typing (cgMLST) and single-nucleotide polymorphism (SNP)-based analyses (13, 16). These improvements make ONT a practical platform for real-time genomic surveillance of bacterial pathogens. ONT has already been successfully used to identify *Salmonella* serotypes, AMR determinants, and resistance structures (17); however, large-scale evaluations of its performance in predicting phenotypic AMR remain limited. Notably, the concordance between genotypic AMR profiles and phenotypic AST results has not been systematically assessed using large collections of *Salmonella* isolates.

In this study, we applied ONT sequencing to a large-scale collection of 1,490 *Salmonella* isolates obtained through nationwide surveillance in Taiwan in 2025. We conducted comprehensive genomic characterization, including the identification of AMR determinants, serotype prediction, multilocus sequence typing (MLST), and plasmid replicon profiling. The primary objective was to evaluate the performance of ONT-based WGS for predicting AMR. Specifically, we assessed the concordance between genotypic AMR profiles and phenotypic AST results. Additionally, we utilized WGS data to track the circulation of major MDR and extensively drug-resistant (XDR) *Salmonella* clones previously reported in Taiwan.

## MATERIALS AND METHODS

### Bacterial isolates

A total of 1,490 *Salmonella* isolates were collected from hospitals across Taiwan through a nationwide surveillance program. Clinical isolates submitted by participating hospitals were initially cultured on Salmonella–Shigella (SS) agar for selective isolation and purification. Presumptive *Salmonella* colonies were identified based on characteristic colony morphology and subsequently subcultured onto tryptic soy agar (TSA) to obtain sufficient biomass for downstream applications. Genomic DNA was extracted from TSA-grown cultures for WGS. In parallel, purified isolates grown on TSA were preserved for long-term storage in tryptic soy broth supplemented with 15% glycerol at −80 °C. For AST, TSA-grown isolates were subcultured onto blood agar plates to ensure optimal growth prior to testing.

### Whole-genome sequencing and analysis

ONT-WGS was performed using the MinION devices (Mk1B or Mk1D) from Oxford Nanopore Technologies (Oxford, United Kingdom). Genomic DNA was extracted from TSA-grown cultures using the EZ2 PowerFecal Pro DNA/RNA Kit (cat. no. 954634; Qiagen, Hilden, Germany), according to the manufacturer’s instructions. Sequencing libraries were prepared using the Rapid Barcoding Kit (SQK-RBK114; Oxford Nanopore Technologies) and sequenced on MinION devices equipped with R10.4.1 flow cells (FLO-MIN114). Raw signal data (POD5 format) were basecalled using Dorado v1.1.1 (Oxford Nanopore Technologies) with the Super Accurate (SUP5.2.0) model to generate FASTQ reads. Long-read assemblies were generated using Flye v2.9.6 (18). Circular contigs were re-oriented using Dnaapler v1.2.0 (19) and subsequently polished using Medaka v2.0.1 (Oxford Nanopore Technologies).

In silico serotype prediction was performed using SISTR v1.1.3 (20), and sequence types (STs) were assigned using mlst v2.32.2 (https://github.com/tseemann/mlst). Plasmid replicon types were identified using PlasmidFinder v2.2.0 (21). AMR determinants, including acquired resistance genes and resistance-associated mutations, were identified using AMRFinderPlus v4.0.3 (22).

### Antimicrobial susceptibility testing

AST was performed using the EUVSEC3 Sensititre broth microdilution MIC panel (TREK Diagnostic Systems Ltd., Thermo Fisher Scientific, East Grinstead, United Kingdom), which includes 15 antimicrobial agents recommended for antimicrobial resistance monitoring in *Salmonella* under the European Union surveillance protocol. Minimum inhibitory concentrations (MICs) were interpreted according to the Clinical and Laboratory Standards Institute (CLSI) guidelines (23) for 14 antimicrobial agents. For tigecycline, interpretive criteria were based on the European Committee on Antimicrobial Susceptibility Testing (EUCAST) recommendations (24). Multidrug-resistant (MDR) isolates were defined as isolates resistant to the antimicrobial agents traditionally used to treat *Salmonella* infections, including ampicillin, chloramphenicol, and trimethoprim-sulfamethoxazole. XDR isolates were defined as MDR isolates that additionally exhibited nonsusceptibility to fluoroquinolones and resistance to third-generation cephalosporins. AziR-XDR isolates were defined as XDR isolates that were also resistant to azithromycin. In cases where phenotypic resistance and AMR determinants-inferred resistance were inconsistent for one or more antimicrobial agents, AST or WGS was repeated to verify.

### Data availability

The whole-genome sequencing data generated in this study have been deposited in the NCBI database under BioProject PRJNA478278. BioSample and SRA accession numbers for the 1,490 isolates are listed in the Supplementary Table S1.

## RESULTS

### Phenotypic and genotypic resistance

Among the 1,490 *Salmonella* isolates, phenotypic AMR varied across antimicrobial classes (Table 1). High resistance rates (>50%) were observed for ampicillin (60.5%), sulfamethoxazole (60.8%), tetracycline (60.3%), and chloramphenicol (53.2%). A substantial proportion of isolates also exhibited ciprofloxacin nonsusceptibility (46.4%) and trimethoprim resistance (45.2%). Moderate resistance rates were noted for cefotaxime (21.5%), tigecycline (20.1%), ceftazidime (19.3%), nalidixic acid (14.8%), and gentamicin (11.7%), whereas a lower resistance rate was observed for azithromycin (4.4%). Resistance to amikacin and meropenem was rare, each detected in a single isolate (0.1%). For colistin, resistance differed markedly by serovar. Colistin resistance was rare (0.8%) in non-*S*. Enteritidis isolates, whereas a high resistance rate (35.4%) was observed in *S*. Enteritidis isolates.

**Table 1.** Comparison of antimicrobial resistance determined by phenotypic antimicrobial susceptibility testing (AST) and inferred from whole-genome sequencing (WGS)–identified resistance determinants among 1,490 *Salmonella* isolates.

| Antimicrobial | Phenotypic resistance |  | Genotypic resistance |  | Resistance determinants |
| --- | --- | --- | --- | --- | --- |
|  | No. isolates | Resistance, % | No. isolates (included *ramAp*) | Resistance, % (included *ramAp*) |  |
| Amikacin | 1 | 0.1 | 1 | 0.1 | *rmtB1* |
| Ampicillin | 902 | 60.5 | 905 | 60.7 | *ampC*, *bla*<sub>CMY</sub>, *bla*<sub>CMY-2</sub>, *bla*<sub>CTX-M</sub>, *bla*<sub>CTX-M-14</sub>, *bla*<sub>CTX-M-15</sub>, *bla*<sub>CTX-M-3</sub>, *bla*<sub>CTX-M-55</sub>, *bla*<sub>CTX-M-65</sub>, *bla*<sub>DHA-1</sub>, *bla*<sub>KPC-2</sub>, *bla*<sub>LAP-2</sub>, *bla*<sub>OXA</sub>, *bla*<sub>OXA-10</sub>, *bla*<sub>PSE</sub>, *bla*<sub>TEM</sub>, *bla*<sub>TEM-1</sub>, *bla*<sub>TEM-135</sub>, *bla*<sub>TEM-176</sub> |
| Azithromycin | 66 | 4.4 | 61 (197) | 4.1 (13.2) | *erm(42)*, *erm(B)*, *mph(A)*, *ramAp* |
| Cefotaxime | 320 | 21.5 | 351 | 23.6 | *ampC*, *bla*<sub>CMY</sub>, *bla*<sub>CMY-2</sub>, *bla*<sub>CTX-M</sub>, *bla*<sub>CTX-M-14</sub>, *bla*<sub>CTX-M-15</sub>, *bla*<sub>CTX-M-3</sub>, *bla*<sub>CTX-M-55</sub>, *bla*<sub>CTX-M-65</sub>, *bla*<sub>DHA-1</sub>, *bla*<sub>KPC-2</sub> |
| Ceftazidime | 288 | 19.3 | 351 | 23.6 | *ampC*, *bla*<sub>CMY</sub>, *bla*<sub>CMY-2</sub>, *bla*<sub>CTX-M</sub>, *bla*<sub>CTX-M-14</sub>, *bla*<sub>CTX-M-15</sub>, *bla*<sub>CTX-M-3</sub>, *bla*<sub>CTX-M-55</sub>, *bla*<sub>CTX-M-65</sub>, *bla*<sub>DHA-1</sub>, *bla*<sub>KPC-2</sub> |
| Chloramphenicol | 793 | 53.2 | 789 (795) | 53.0 (53.4) | *catA1*, *catA2*, *cmlA1*, *cmlA5*, *floR*, *ramAp* |
| Ciprofloxacin<sup>NS</sup> | 692 | 46.4 | 696 (711) | 46.7 (47.7) | *aac(6')-Ib-cr*, *gyrA_D87G*, *gyrA_D87N*, *gyrA_D87Y*, *gyrA_S83F*, *gyrA_S83Y*, *parC_S80I*, *qepA8*, *qnrA1*, *qnrB*, *qnrB6*, *qnrD1*, *qnrS*, *qnrS1*, *qnrS13*, *ramAp* |
| Colistin (all but *S. Enteritidis*, N=1233) | 10 | 0.8 | 9 | 0.7 | *mcr-1.1* |
| Colistin (*S. Enteritidis*, N=257) | 91 | 35.4 | 1 | 0.1 | *mcr-1.1* |
| Gentamicin | 174 | 11.7 | 174 | 11.7 | *aac(3)-IId*, *aac(3)-Ile*, *aac(3)-IVa*, *rmtB1* |
| Meropenem | 1 | 0.1 | 1 | 0.1 | *bla*<sub>KPC-2</sub> |
| Nalidixic acid | 221 | 14.8 | 118 (254) | 7.9 (17.0) | *gyrA_D87G*, *gyrA_D87N*, *gyrA_D87Y*, *gyrA_S83F*, *gyrA_S83Y*, *parC_S80I*, *ramAp* |
| Sulfamethoxazole | 906 | 60.8 | 907 (914) | 60.9 (61.3) | *ramAp*, *sul1*, *sul2*, *sul3* |
| Tetracycline | 899 | 60.3 | 904 (907) | 60.7 (60.9) | *ramAp*, *tet(A)*, *tet(B)*, *tet(D)*, *tet(G)*, *tet(M)* |
| Tigecycline | 299 | 20.1 | 0 (139) | 0 (9.3) | *ramAp* |
| Trimethoprim | 673 | 45.2 | 674 (677) | 45.2 (45.4) | *dfrA1*, *dfrA12*, *dfrA14*, *dfrA17*, *dfrA23*, *dfrA25*, *dfrA27*, *dfrA5*, *dfrA51*, *dfrA7*, *ramAp* |
| Clindamycin | NT | NT | 322 | 21.6 | *erm(42)*, *erm(B)*, *lnu(F)* |
| Erythromycin | NT | NT | 93 | 6.2 | *erm(42), erm(B), estT, mph(A)* |
| Fosfomicin | NT | NT | 203 | 13.6 | *fosA3, fosA7, fosA7.2, fosA7.3, fosA7.4* |
| Hygromycin B | NT | NT | 31 | 2.1 | *aph(4)-Ia* |
| Kanamycin | NT | NT | 333 | 22.3 | *aac(6'), aac(6')-Ib-cr, aph(3')-Ia, aph(3')-Iia* |
| Rifampin | NT | NT | 166 | 11.1 | *arr-2, arr-3* |
| Streptomycin | NT | NT | 896 | 60.1 | *aadA1, aadA16, aadA2, aadA22, aadA7, aph(3'')-Ib, aph(6)-Ic, aph(6)-Id* |

\*Genotypic resistance was inferred from acquired resistance genes and resistance-associated mutations (AMR determinants). Values in parentheses indicate the number and percentage of isolates harboring AMR determinants, *ramAp* alone, or both. NS, non-susceptible; NT, not tested.

Genotypic resistance inferred from WGS-identified AMR determinants showed trends similar to those of phenotypic resistance for most antimicrobials (Table 1). Concordant or near-concordant resistance proportions (excluding the effect of *ramAp*) were observed for ampicillin (60.5% vs. 60.7%), azithromycin (4.4% vs. 4.1%), chloramphenicol (53.2% vs. 53.0%), ciprofloxacin nonsusceptibility (46.2% vs. 46.7%), colistin in non-*S*. Enteritidis isolates (0.8% vs. 0.7%), gentamicin (11.7% vs. 11.7%), meropenem (0.1% vs. 0.1%), sulfamethoxazole (60.8% vs. 60.9%), tetracycline (60.3% vs. 60.7%), and trimethoprim (45.2% vs. 45.2%).

Marked discrepancies between phenotypic and genotypic resistance were observed for selected antimicrobials. For tigecycline, no resistance was predicted based on known AMR determinants, whereas 20.1% of isolates were phenotypically resistant under EUCAST criteria (S ≤0.5; R >0.5 mg/L). Similarly, for nalidixic acid, genotypic resistance excluding the effect of *ramAp* (7.9%) was substantially lower than the observed phenotypic resistance (14.8%). In contrast, genotypic resistance exceeded phenotypic resistance for cefotaxime (23.6% vs. 21.5%) and ceftazidime (23.6% vs. 19.3%).

For colistin, a pronounced discrepancy was observed in *S*. Enteritidis isolates, where 35.4% (91/257) were phenotypically resistant, whereas only 0.1% (1/257) carried *mcr-1*.*1*. In contrast, among non-*S*. Enteritidis isolates, both phenotypic and genotypic resistance rates were low (0.8% and 0.7%, respectively). Consistent with this pattern, among isolates lacking known colistin resistance determinants, *S*. Enteritidis isolates exhibited a shift toward higher colistin MIC values, whereas non-*S*. Enteritidis isolates were predominantly distributed at lower MIC levels (Supplementary Fig. S1).

Incorporation of *ramAp* markedly affected genotype-based resistance prediction for several antimicrobials (Table 1). *ramAp*, a plasmid-borne *ramA* homolog, has been shown to elevate expression of the AcrAB-TolC efflux system and to elevate MICs to multiple antimicrobials, including azithromycin, cefoxitin, chloramphenicol, ciprofloxacin, nalidixic acid, sulfamethoxazole, tetracycline, tigecycline, and trimethoprim (25). Incorporating *ramAp* into resistance predictions led to a significant increase in the proportion of isolates classified as genotypically resistant, especially for azithromycin (from 4.1% to 13.2%), nalidixic acid (from 7.9% to 17.0%), and tigecycline (from 0% to 9.3%). In contrast, only minor increases were observed for chloramphenicol (from 53.0% to 53.4%), ciprofloxacin nonsusceptibility (from 46.7% to 47.7%), sulfamethoxazole (from 60.9% to 61.3%), tetracycline (from 60.7% to 60.9%), and trimethoprim (from 45.2% to 45.4%). In *ramAp*-positive isolates, resistance to chloramphenicol, ciprofloxacin, sulfamethoxazole, tetracycline, and trimethoprim was already largely attributable to other established AMR determinants. In contrast, known AMR determinants for azithromycin, nalidixic acid, and tigecycline were relatively uncommon. As a result, the contribution of *ramAp*-associated MIC elevation was more evident for these antimicrobials.

Although several antimicrobials were not included in the AST panel, their resistance rates could be predicted based on AMR determinants identified from WGS data. These antimicrobials included streptomycin (60.1%), kanamycin (22.3%), clindamycin (21.6%), fosfomycin (13.6%), rifampin (11.1%), erythromycin (6.2%), and hygromycin B (2.1%) (Table 1).

### Genotype-phenotype discordance and contributing factors

Genotype–phenotype resistance discordance among the 1,490 *Salmonella* isolates (Table 2) could be categorized into four major types: (i) breakpoint-dependent classification, (ii) reduced or absent phenotypic expression of resistance genes, (iii) MIC modulation by *ramAp*, and (iv) absence of known AMR determinants.

**Table 2.** Categories of genotype–phenotype discordance and contributing factors affecting antimicrobial susceptibility among 1,490 *Salmonella isolates*.

| Category | Antimicrobial | No. isolates | Description of discrepancy | Reference |
| --- | --- | --- | --- | --- |
| Breakpoint-dependent classification | Cefotaxime | 31 | *bla*<sub>DHA-1</sub>- and *bla*<sub>CTX-M-65</sub>-carrying isolates classified as susceptible or intermediate under CLSI criteria | Table S2_FOT |
|  | Ceftazidime | 63 | *ampC*-, *bla*<sub>CMY-2</sub>-, *bla*<sub>CTX-M-3</sub>-, *bla*<sub>CTX-M-65</sub>-, and *bla*<sub>DHA-1</sub>-carrying isolates classified as susceptible or intermediate under CLSI criteria | Table S2_TAZ |
| Reduced or absent phenotypic expression of resistance genes | Ampicillin | 3 | *bla*<sub>CTX-M</sub> and *bla*<sub>TEM-1</sub>-carrying isolates exhibited ampicillin MICs within the intermediate range | Table S2_AMP |
|  | Ciprofloxacin | 4 | Isolates carrying *qnrS1* only exhibited a ciprofloxacin MIC of 0.06 mg/L, below the resistance breakpoint | Table S2_CIP |
|  | Sulfamethoxazole | 1 | Isolate carrying *sul1* exhibited a sulfamethoxazole MIC of 16 mg/L, below the resistance breakpoint | Table S2_SUL |
|  | Tetracycline | 5 | Isolates carrying *tet(M)* only exhibited tetracycline MICs of 2–4 mg/L, below the resistance breakpoint | Table S2_TET |
|  | Trimethoprim | 1 | Isolate carrying *dfrA51* exhibited a trimethoprim MIC of 0.25 mg/L, below the resistance breakpoint | Table S2_TMP |
| MIC modulation by *ramAp* | Azithromycin | 5 | Isolates carrying *ramAp* only exhibited elevated MICs reaching the resistance breakpoint | Table S2_AZI |
|  | Chloramphenicol | 4 | Isolates carrying *ramAp* only exhibited elevated MICs reaching the resistance breakpoint | Table S2_CHL |
|  | Nalidixic acid | 100 | Isolates carrying *ramAp* only exhibited elevated MICs reaching the resistance breakpoint | Table S2_NAL |
|  | Tigecycline | 135 | Isolates carrying *ramAp* only exhibited elevated MICs reaching the resistance breakpoint | Table S2_TGE1 |
| Absence of known AMR determinants | Nalidixic acid | 3 | Isolates lacking known resistance determinants exhibited a nalidixic acid MIC at the resistance breakpoint (32 mg/L) | Table S2_NAL |
|  | Tigecycline | 164 | Isolates lacking known tigecycline resistance determinants nevertheless exhibited tigecycline MICs at or above the resistance breakpoint | Table S2_TGE1 |
|  | Colistin | 91 | Ninety *S. Enteritidis* isolates and one *S. Chincol* isolate lacked known colistin resistance determinants despite exhibiting phenotypic resistance | Table S2_COL |

The first category comprised breakpoint-dependent classification for cefotaxime and ceftazidime. Using CLSI interpretive criteria for cefotaxime (S ≤1, I = 2, R ≥4 mg/L), 31 isolates carrying *bla*_CTX-M-65_ or *bla*_DHA-1_ were classified as susceptible or intermediate (Table S2_FOT). Similarly, 63 isolates carrying β-lactamase genes, including *ampC, bla*_CMY-2_, *bla*_CTX-M-3_, *bla*_CTX-M-65_, and *bla*_DHA-1_, were categorized as susceptible or intermediate for ceftazidime under CLSI criteria (S ≤4, I = 8, R ≥16 mg/L) (Table S2_TAZ).

The second category, reduced or absent phenotypic expression of resistance genes, was observed for ampicillin, ciprofloxacin, sulfamethoxazole, tetracycline, and trimethoprim. In these cases, isolates carrying known AMR determinants, including *bla*_CTX-M_, *bla*_TEM-1_, *qnrS1, sul1, tet(M)*, and *dfrA51*, remained phenotypically susceptible or exhibited only intermediate resistance, showing that some isolates carrying these determinants exhibited MICs below the clinical resistance breakpoints (Tables S2_AMP, S2_CIP, S2_SUL, S2_TET, and S2_TMP).

The third category involved efflux-associated MIC elevation linked to *ramAp*. For azithromycin and chloramphenicol, five isolates carrying *ramAp* alone exhibited MICs at or above the resistance breakpoint (Tables S2_AZI and S2_CHL). A similar pattern was observed for nalidixic acid, in which 100 isolates carrying *ramAp* alone exhibited MICs reaching the resistance breakpoint (Table S2_NAL). For tigecycline, 135 isolates carrying *ramAp* alone exhibited MICs at or above the EUCAST resistance breakpoint (Table S2_TGE1).

The fourth category involved phenotypic resistance in the absence of known AMR determinants. For nalidixic acid, three isolates lacking known AMR determinants exhibited MICs at the resistance breakpoint (32 mg/L) (Table S2_NAL). Similarly, 164 isolates lacking known tigecycline resistance determinants nevertheless exhibited tigecycline MICs at or above the EUCAST resistance breakpoint (Table S2_TGE1). For colistin, 91 isolates lacking known AMR determinants were phenotypically resistant, including 90 *S*. Enteritidis isolates and one *S*. Chincol isolate (Table S2_COL).

### Serovar distribution and associated antimicrobial resistance patterns

A total of 64 major serovars were identified among 1,490 isolates, with *S*. I 1,4,[5],12:i:-(26.0%) being the most prevalent, followed by *S*. Enteritidis (17.2%) and *S*. Agona (10.1%) (Table 3). Other commonly detected serovars included *S*. Anatum (6.0%), *S*. Newport (4.4%), *S*. Typhimurium (4.4%), *S*. Kentucky (4.2%), *S*. Goldcoast (2.8%), *S*. Stanley (2.8%), and *S*. Infantis (1.9%).

**Table 3.**
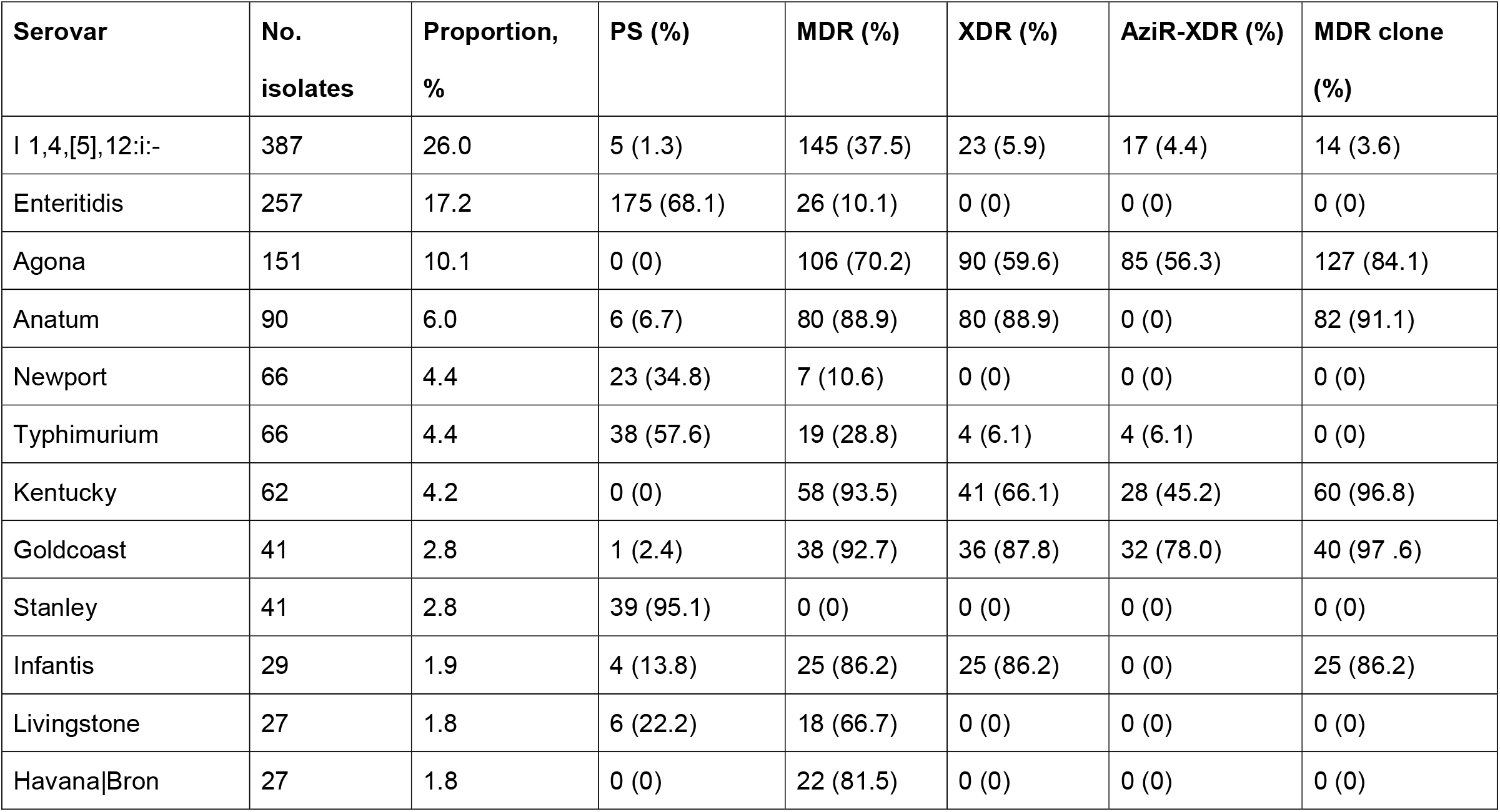
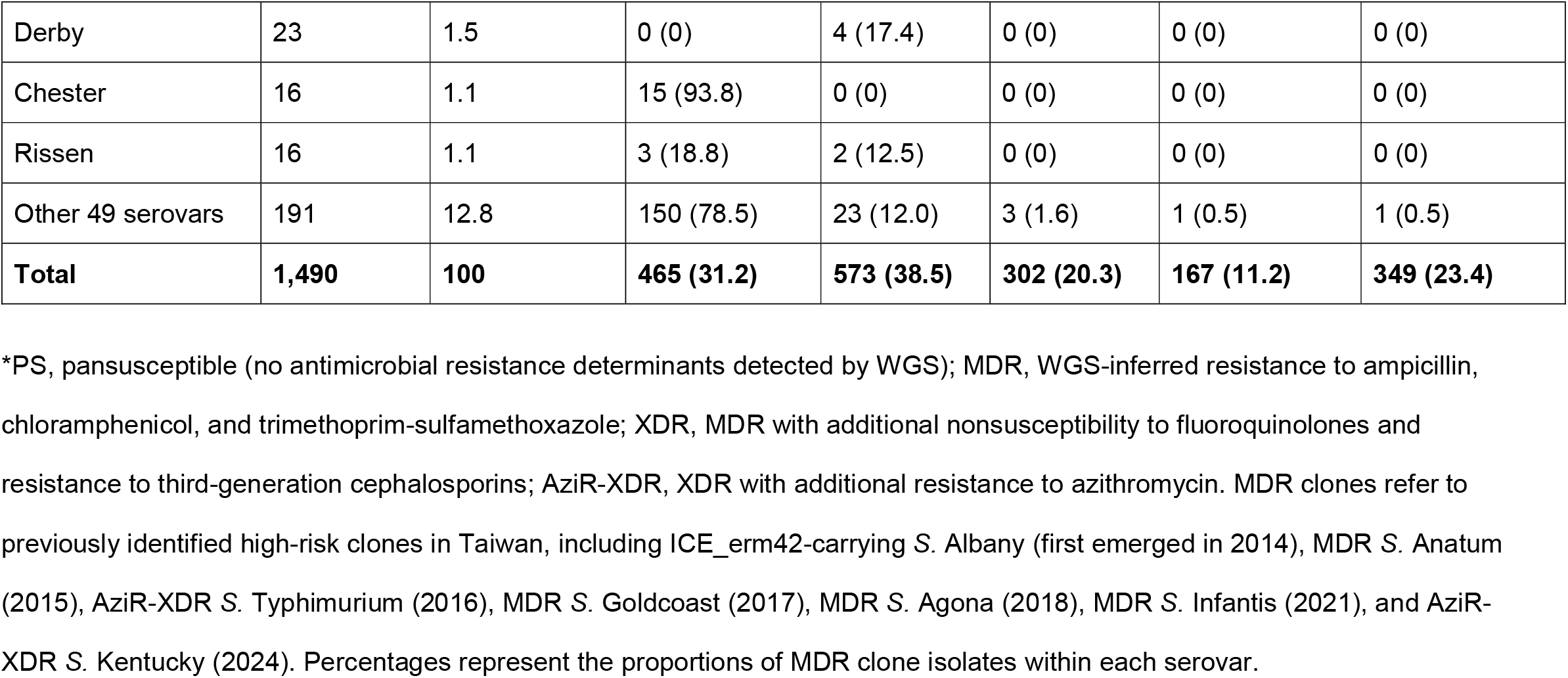
Distribution of Salmonella serovars and antimicrobial resistance profiles among 1,490 isolates collected in Taiwan, 2025.

Antimicrobial resistance profiles varied markedly across serovars, as predicted based on AMR determinants identified through WGS. Among the 15 most common serovars, a high proportion of pansusceptible isolates was observed in *S*. Stanley (95.1%), *S*. Chester (93.8%), *S*. Enteritidis (68.1%), and S. Typhimurium (57.6%), whereas MDR was highly prevalent in *S*. Kentucky (93.5%), *S*. Goldcoast (92.7%), *S*. Anatum (88.9%), *S*. Infantis (86.2%), *S*. Havana (81.5%), and *S*. Agona (70.2%). XDR phenotypes were most frequently observed in *S*. Anatum (88.9%), *S*. Goldcoast (87.8%), *S*. Infantis (86.2%), *S*. Kentucky (66.1%), and *S*. Agona (59.6%). AziR-XDR isolates were predominantly identified in *S*. Goldcoast (78.0%), *S*. Agona (56.3%), *S*. Kentucky (45.2%), with lower proportions observed in *S*. Typhimurium (6.1%), and *S*. I 1,4,[5],12:i:-(4.4%). Overall, 31.2% of isolates were pansusceptible, 38.5% were MDR, 20.3% were XDR, and 11.2% were AziR-XDR.

### Epidemiological distribution and persistence of major MDR clones

Previously identified MDR clones from long-term surveillance (2004–2024), including ICE_erm42-carrying S. Albany (26), MDR *S*. Anatum (6), *S*. Goldcoast (5), *S*. Agona (8), *S*. Infantis (9), and *S*. Typhimurium (7), as well as MDR *S*. Kentucky newly identified in 2025 (10), were detected in the 2025 isolate collection. Based on WGS-derived phylogenetic analysis and resistance profiles, these MDR clones constituted high proportions within their respective serovars, including *S*. Goldcoast (97.6%), *S*. Kentucky (96.8%), *S*. Anatum (91.1%), *S*. Infantis (86.2%), and *S*. Agona (84.1%). One ICE_erm42-carrying S. Albany was also detected in this isolate collection. In total, these MDR clones accounted for 23.4% of all isolates (Table 3).

### Association of *ramAp* with MIC elevation

In this study, the effects of the plasmid-borne *ramAp* were evaluated for azithromycin, nalidixic acid, and tigecycline, as sufficient isolates carrying *ramAp* alone were available for analysis. MIC distributions revealed consistent shifts toward higher MIC values for isolates carrying *ramAp* alone, compared to wild-type isolates. For azithromycin and tigecycline, MIC distributions shifted by approximately 4-fold, while nalidixic acid exhibited a more pronounced shift of ≥8-fold (Supplementary Figures S2–S4).

Thirty-five *S*. Enteritidis isolates carrying *ramAp* were excluded from the antimicrobial mapping analysis because they lacked an upstream promoter region, indicating that *ramAp* was unlikely to be expressed in these isolates.

## DISCUSSION

This study provides a comprehensive evaluation of genotype–phenotype resistance in 1,490 *Salmonella* isolates using ONT-based WGS data in a nationwide surveillance setting. Overall, genotypic resistance predictions demonstrated high concordance with phenotypic AST results across most antimicrobial classes, supporting the utility of ONT-WGS as a practical tool for large-scale AMR identification and surveillance (Table 1). Concordance was particularly high for several clinically important antimicrobials, including ampicillin, azithromycin, chloramphenicol, ciprofloxacin, gentamicin, meropenem, sulfamethoxazole, tetracycline, and trimethoprim, for which phenotypic and genotypic resistance rates differed by ≤0.5% (Table 1). In addition to predicting resistance phenotypes, WGS data also provided detailed information on AMR determinants, resistance-associated mutations, and the genetic contexts of resistance genes that could not be obtained from conventional AST alone. However, notable genotype–phenotype discrepancies remained for selected antimicrobial classes, particularly third-generation cephalosporins, tigecycline, and colistin. These discordances were not random but were associated with several biologically and methodologically distinct mechanisms, including breakpoint-dependent classification, reduced or absent phenotypic expression of resistance genes, MIC modulation by *ramAp*, and the absence of known AMR determinants (Table 2).

Breakpoint-dependent classification contributed to discordance for β-lactam antimicrobials, particularly cefotaxime and ceftazidime. A subset of isolates carrying β-lactamase genes, including *bla*_CTX-M-65_ and *bla*_DHA-1_, were classified as susceptible or intermediate under CLSI criteria. This discordance was most evident for cefotaxime, where among 117 *bla*_DHA-1_-carrying isolates, 30 (25.6%) exhibited MICs of 1–2 mg/L and therefore fell below the resistance breakpoint (Table S2_FOT). A similar pattern was observed for ceftazidime, where 25 of 26 *bla*_CTX-M-65_-carrying isolates exhibited MICs of 2–4 mg/L and were categorized as susceptible (Table S2_TAZ). In addition, 33 (28.2%) of *bla*_DHA-1_-carying isolates were classified as non-resistant. These findings suggest that the current CLSI breakpoints may not reliably distinguish wild-type from β-lactamase–producing isolates in this setting.

Importantly, such discordance has been associated with adverse clinical outcomes. Previous studies have demonstrated that infections caused by ESBL-producing organisms may result in treatment failure when treated with cephalosporins despite in vitro susceptibility. In a landmark study, Paterson et al. reported failure rates of 54% among patients infected with ESBL-producing organisms classified as susceptible and 100% among those with intermediate MICs (27). Pharmacokinetic and pharmacodynamic analyses further support these observations, showing that as MIC values approach clinical breakpoints, the probability of achieving pharmacodynamic targets (e.g., %T > MIC) declines substantially, even for isolates categorized as susceptible, thereby compromising therapeutic efficacy (28). Our data highlight that the current interpretive criteria (clinical breakpoints) for cefotaxime and ceftazidime do not effectively distinguish wild-type from non-wild-type populations. These isolates may be misclassified as non-resistant despite carrying AMR determinants, increasing the risk of treatment failure. In contrast, the use of epidemiological cutoff values (ECOFFs) as a complementary framework improves the differentiation between wild-type and non-wild-type populations, enhancing concordance between genotypic and phenotypic resistance. In our data, the MIC distributions suggest tentative ECOFFs of >0.5 mg/L (≥1 mg/L) for cefotaxime (Table S2_FOT) and >1 mg/L (≥2 mg/L) for ceftazidime (Table S2_TAZ), which more effectively separate wild-type isolates from those carrying β-lactamase determinants and reduce the risk of misclassification.

The plasmid-borne regulator *ramAp* plays a crucial role in complicating the interpretation of genotype–phenotype resistance. Unlike traditional resistance genes, *ramAp* does not directly cause high-level resistance. Instead, it increases MICs by activating multidrug efflux systems, which lowers intracellular drug concentrations (25). Our data indicate that *ramAp* was associated with approximately 4-fold increases in MICs for tigecycline and azithromycin (Figure S2 and Figure S4) and ≥8-fold increases for nalidixic acid (Figure S3). Despite these effects, most *ramAp*-carrying isolates remained below clinical breakpoints and were therefore categorized as susceptible or intermediate. This complicates interpretation because isolates with elevated MIC values may still be classified as susceptible under current clinical breakpoints, potentially leading to misclassification of their resistance profiles. From a clinical perspective, such shifts may reduce pharmacodynamic margins, thereby compromising treatment efficacy in certain cases. These findings highlight a critical limitation of binary susceptibility interpretation, as isolates categorized as susceptible may still harbor MIC-modulating mechanisms like *ramAp*, which are not captured by conventional genotype-based predictions. This underscores the need to better account for *ramAp* in clinical decision-making.

Substantial genotype–phenotype discordance was observed for tigecycline and colistin (Table 1). For tigecycline, 20.1% of isolates were phenotypically resistant under EUCAST criteria, yet no known high-level resistance genes, such as *tet(X)*, were detected (29, 30). This finding is consistent with previous studies suggesting that tigecycline resistance in *Salmonella* is often mediated by non-classical mechanisms, including efflux overexpression and ribosomal protection-associated changes (31, 32). Of the 299 tigecycline-resistant isolates, 139 showed resistance linked to *ramAp*, which contributed to an approximately fourfold increase in MICs (Figure S4). However, the remaining 164 isolates had resistance mechanisms that are currently unknown, highlighting the need for further investigation into additional tigecycline resistance determinants (Table S2_TGE2). These findings suggest that the mechanisms responsible for tigecycline resistance may involve gradual MIC elevation through multiple, less-characterized factors, rather than distinct, well-defined resistance genes.

The most striking genotype–phenotype discordance was observed for colistin in *S*. Enteritidis. A high rate of phenotypic resistance was identified, with 35.4% (91/257) of isolates exceeding the clinical breakpoint, yet only one isolate carried a known AMR determinant (*mcr-1*.*1*). Among isolates lacking known colistin resistance determinants, 35.2% (90/256) exhibited MICs at or above the resistance threshold, accompanied by a right-shifted MIC distribution (Figure S1). This pattern aligns with previous reports from Taiwan and China, where high rates of colistin resistance in *S*. Enteritidis (72.4% and 83.9%, respectively) were not explained by *mcr* genes or canonical chromosomal mutations (4, 33). Notably, mutations in known regulatory systems (e.g., *pmrAB*) are rarely detected in these resistant isolates, suggesting that the observed phenotype is not attributable to currently recognized resistance mechanisms. Studies by Agersø et al.(34) and Ricci et al. (35) support the notion that elevated colistin MICs in *S*. Enteritidis and other O:1,9,12 serovars, such as *S*. Dublin, represent a serovar-associated intrinsic resistance phenotype. This resistance is likely linked to differences in O-antigen structure, which may prevent colistin from reaching its target.

Another category of discordance was reduced or absent phenotypic expression of resistance genes, although such cases were infrequent in this study (Table 2). Specifically, five isolates carrying *tet(M)* exhibited low tetracycline MICs (2–4 mg/L) and were phenotypically susceptible despite carrying intact gene sequences and upstream promoter regions (Table S2_TET). Similarly, isolates carrying other resistance genes, such as *qnrS1* for ciprofloxacin and *sul1* for sulfamethoxazole, showed low MICs below resistance breakpoints (Table S2_CIP, Table S2_SUL). These observations suggest that these ARGs might either be insufficiently expressed or not expressed at all.

In addition, some discordant cases were attributed to gene truncation or regulatory disruption. For example, 9 *cmlA1*-carrying *S*. Rissen isolates, initially annotated as having complete genes based on the default coverage threshold (≥90%), were later found to have partial sequences (coverage 90.7%) upon re-analysis due to genotype–phenotype inconsistency. Similarly, *aac(3)-Iva* in two isolates exhibited an IS26-mediated promoter interruption, explaining the absence of the gentamicin resistance phenotype. These findings underscore the importance of considering both gene integrity and regulatory context in genomic analyses.

In addition to AMR profiling, this study underscores the value of WGS-based surveillance in monitoring the epidemiological dynamics of previously identified high-risk MDR clones in Taiwan. As shown in Table 3, serovars associated with these MDR clones, including *S*. Anatum (first identified in 2015), *S*. Goldcoast (2017), *S*. Agona (2018), *S*. Infantis (2021), and *S*. Kentucky (2024), remained among the most prevalent serovars in 2025. Notably, MDR clones continued to represent the dominant proportion within their respective serovars, comprising 91.1% of *S*. Anatum, 97.6% of *S*. Goldcoast, 84.1% of *S*. Agona, 86.2% of *S*. Infantis, and 96.8% of *S*. Kentucky. These findings indicate the ongoing circulation and persistence of established MDR clones over time, rather than the sporadic or transient emergence of new clones. The high proportion of MDR clones within these serovars suggests that clonal expansion remains a key driver of antimicrobial resistance in *Salmonella* populations in Taiwan. Collectively, these results highlight the importance of integrating WGS into routine surveillance, not only for resistance prediction but also for tracking the long-term persistence and spread of high-risk clones.

In conclusion, this study demonstrates that ONT-WGS provides a highly reliable method for AMR prediction in *Salmonella*, with genotype–phenotype concordance observed in the vast majority of isolates. However, notable discordance was found for specific antimicrobial classes, such as tigecycline and colistin, which were linked to non-classical resistance mechanisms, MIC modulation by elements like *ramAp*, and breakpoint-dependent classification. Despite these exceptions, ONT-WGS provides faster and more comprehensive AMR data, which could complement or even replace conventional AST in the future, offering valuable insights for clinical therapy and epidemiological surveillance.

## SUPPLEMENTAL MATERIAL

Supplemental material is available online only.

Supplemental figures, PDF file, 0.37 MB.

Supplemental tables, XLSX file, 0.62 MB.

## ACKNOWLEDGMENTS

This work was supported by the Ministry of Health and Welfare, Taiwan, under grants MOHW115-CDC-C-305-114304, MOHW113-CDC-C-315-114110, MOHW114-CDC-C-315-124112, and MOHW115-CDC-C-305-134118.

